# Rapid assessment of susceptibility of bacteria and erythrocytes to antimicrobial peptides by single-cell impedance cytometry

**DOI:** 10.1101/2022.11.04.515154

**Authors:** Cassandra Troiano, Adele De Ninno, Bruno Casciaro, Francesco Riccitelli, Yoonkyung Park, Renato Massoud, Maria Luisa Mangoni, Paolo Bisegna, Lorenzo Stella, Federica Caselli

## Abstract

Antimicrobial peptides (AMPs) represent a promising class of compounds to fight antibiotic-resistant infections. In most cases, they kill bacteria by making their membrane permeable and therefore exhibit low propensity to induce bacterial resistance. In addition, they are often selective, killing bacteria at concentrations lower than those at which they are toxic to the host. However, clinical applications of AMPs are hindered by a limited understanding of their interactions with bacteria and human cells. Standard susceptibility testing methods are based on the analysis of the growth of a bacterial population and therefore require several hours. Moreover, different assays are required to assess the toxicity to host cells. In this work, we propose the use of microfluidic impedance cytometry to explore the action of AMPs on both bacteria and host cells, in a rapid manner and with single-cell resolution. We show that the electrical signatures of *Bacillus megaterium (B. megaterium)* cells and human red blood cells (RBCs) reflect the action of a representative antimicrobial peptide, DNS-PMAP23. In particular, the impedance phase at high frequency (e.g., 11 or 20 MHz) is a reliable label-free metric for monitoring DNS-PMAP23 bactericidal activity and toxicity to RBCs. The impedance-based characterization is validated by comparison with standard antibacterial activity assays and absorbance-based hemolytic activity assays. Furthermore, we demonstrate the applicability of the technique to a mixed sample of *B. megaterium* cells and RBCs, which paves the way to study AMP selectivity for bacterial versus eukaryotic cells in presence of both cell types.

## 1. Introduction

Many bacterial strains are currently resistant to several, or even all, the available antibiotics. Bacterial resistance to antimicrobial drugs (AMR, for antimicrobial resistance) is a major threat to human health and has been termed an “overlooked pandemic” (Laxminarayan, 2022). More than one million deaths are currently directly attributable to AMR, a value that ranks behind only COVID-19 and tuberculosis in terms of global deaths from an infection (Murray et al., 2022). In addition, the declining efficacy of existing antibiotics is endangering many essential procedures in modern medicine (including surgery, chemotherapy, organ transplantation, etc.) that require effective antimicrobial drugs (Teillant et al., 2015). The problem of AMR is exacerbated by the lack of development of new antibiotics: the last entirely original class of antibiotics was discovered in the late 1980s (Plackett, 2020).

Antimicrobial peptides (AMPs), sometimes also called host defense peptides, are a particularly promising class of molecules to fight AMR (Magana et al., 2020). They are natural molecules produced by all organisms, including humans, as a first line of defense against invading pathogens (Lazzaro et al., 2020). They have a broad spectrum of activity, low toxicity against host cells, and usually kill bacteria in a few minutes, by making their membranes permeable (Matsuzaki, 2019; Roversi et al., 2014). Due to this mechanism of action, the development of resistance against AMPs is particularly difficult (Spohn et al., 2019).

Antimicrobial susceptibility testing (AST), which assesses the susceptibility of pathogens to antimicrobial drugs, is a key factor in the treatment of bacterial infections and in the fight against antibiotic resistance. AST can allow the prescription of appropriate drugs and, therefore, a reduction in the use of broad-spectrum antibiotics. However, current methods require overnight incubation, while a rapid response is critical in the effective treatment of infections. Therefore, there is an urgent need for faster AST methods (Madhusoodanan, 2021; van Belkum et al., 2020). Also in the specific case of AMPs, researchers agree on the necessity of improving AST methods (Meurer et al., 2021). For instance, selectivity for bacterial versus eukaryotic cells is an essential characteristic of AMPs, in view of therapeutic applications. This property arises from the different compositions of the membranes of the two cell types (Bobone and Stella, 2019). However, some of us recently showed that the standard assays used to assess selectivity (separate experiments on human and bacterial cells, at fixed cell densities) are not representative of the real conditions encountered *in vivo* (Loffredo et al., 2021; Savini et al., 2017). The development of approaches where antibiotic activity and toxicity are assessed in the presence of both cell types is essential (Starr et al., 2016). Another limit of current AST methods is that measurements on bacterial populations are not suited to investigate the presence of a small number of persister or viable but non-culturable cells, which are crucial in preventing the eradication of the infection by antimicrobials. Cell-to-cell differences in drug response within a clonal bacterial population have also been reported in the case of AMPs (Cama and Pagliara, 2021; Jepson et al., 2016; Loffredo et al., 2021; Semeraro et al., 2022; Snoussi et al., 2018). Single-cell techniques are required to address cell heterogeneity (Bamford et al., 2017).

Microfluidic impedance cytometry is particularly suited to address the needs of short analysis times and of information at the single-cell level. Its merits with respect to current state-of-the-art AST methods are comprehensively discussed in (Spencer et al., 2020). The technique measures the electrical phenotype of individual biological cells and has been applied to mammalian cells, human pathogens, yeast cells, and plant cells (Daguerre et al., 2020; de Bruijn et al., 2021; Gökçe et al., 2021; Honrado et al., 2021b; Kruit et al., 2022; Petchakup et al., 2018; Tang et al., 2021; Wang et al., 2022). The sensitivity of the technique to alterations of cell size, membrane, and interior composition makes it particularly suitable for cell viability applications (De Ninno et al., 2020; Honrado et al., 2022, 2021a; Zhong et al., 2021). David et al. performed impedance-based viability analysis of *Bacillus megaterium* (*B. megaterium*) with cells at different growth stages and heat-inactivated cells (David et al., 2012). Bertelsen et al. showed that the impedance response of *Escherichia coli* (*E. coli*) depends on its viability state, but the specific response depends on the inactivation method (ethanol, heat and autoclavation) (Bertelsen et al., 2020). Increase in *E. coli* cell volume induced by treatment with Mecillinam could also be detected (Tang et al., 2022). Spencer et al. used microfluidic impedance cytometry to test bacterial susceptibility to traditional antibiotics and showed that the measured electrical characteristics reflect the phenotypic response of the bacteria to the mode of action of a particular antibiotic (Spencer et al., 2020). They also tested the activity of the bacterial, cyclic lipopeptide colistin, which is membrane active. However, to the best of our knowledge, no impedance data are available on the activity of gene-encoded, linear AMPs, belonging to the innate immune system of multicellular organisms, nor on their toxicity towards host cells.

In this paper we present the use of microfluidic impedance cytometry to investigate the activity of AMPs on both bacteria and human cells. We selected the porcine cathelicidin PMAP-23 as a representative example of AMPs. PMAP-23 is a 23 residue, linear, amphipathic peptide, produced by pig myeloid cells (Zanetti et al., 1994), endowed with antibacterial (Zanetti et al., 1994), antifungal (Lee et al., 2001), and antinematodal activities (Park et al., 2004). PMAP-23 kills bacteria by perturbing the permeability of their membranes. It forms pores according to the so-called “carpet” mechanism (Bocchinfuso et al., 2009; Orioni et al., 2009; Roversi et al., 2014), where peptides accumulate on the membrane surface, perturbing its surface tension and causing the formation of defects once a threshold of bound peptide molecules is reached. After the disruption of bacterial membranes, PMAP-23 binds with high affinity to intracellular components (Savini et al., 2020). In recent years, we used a fluorescently labeled analog of PMAP-23 (indicated as DNS-PMAP23, due to the presence of a dansyl label at the N-terminus) to quantitatively characterize its interaction with bacterial and human cells (Roversi et al., 2014; Savini et al., 2020, 2017). For this reason, DNS-PMAP23 was selected for the present study, too. We have previously shown that bacterial killing and hemolysis by DNS-PMAP23 are fast, being completed in less than 15 minutes (Savini et al., 2017). In the present study, the bactericidal activity of DNS-PMAP23 was tested on *B. megaterium* cells, which are commonly used as Gram-positive bacterial model organisms (David et al., 2012). The hemolytic activity of the peptide on human red blood cells (RBCs) form healthy donors was also investigated, as a measure of DNS-PMAP23 cytotoxicity.

The overall experimental protocol is illustrated in **Figure 1** (see **Section 2** for details). Bacterial and RBC samples at different peptide concentrations were prepared (**Fig. 1(a)**) and characterized with reference approaches – a CFU counting assay for the bacterial samples (**Fig. 1(b)**) and an absorbance-based hemolysis assay for the RBC samples (**Fig. 1(c)**) – in parallel to microfluidic impedance cytometry analysis (**Fig. 1(d)**). The results (**Section 3**) show that impedance-based metrics can be used as indicators of AMP-induced cell alterations. Specifically, the impedance phase at high frequency (e.g., 11 or 20 MHz) turns out to be a reliable label-free metric for monitoring AMPs bactericidal activity and toxicity to host cells. Furthermore, a proof-of-concept experiment involving a mixture of *B. megaterium* and RBCs, either incubated with the peptide or untreated, was performed. The simultaneous analysis of bacteria and host cells is critical to study peptide selectivity under realistic conditions. To the best of our knowledge, this is the first time that impedance cytometry is used to characterize a mixture of bacteria and human cells. Overall, these results support the use of microfluidic impedance cytometry for the selection of the most effective AMPs exhibiting maximum activity and minimum toxicity in the presence of mixed cell populations.

**Fig. 1.**
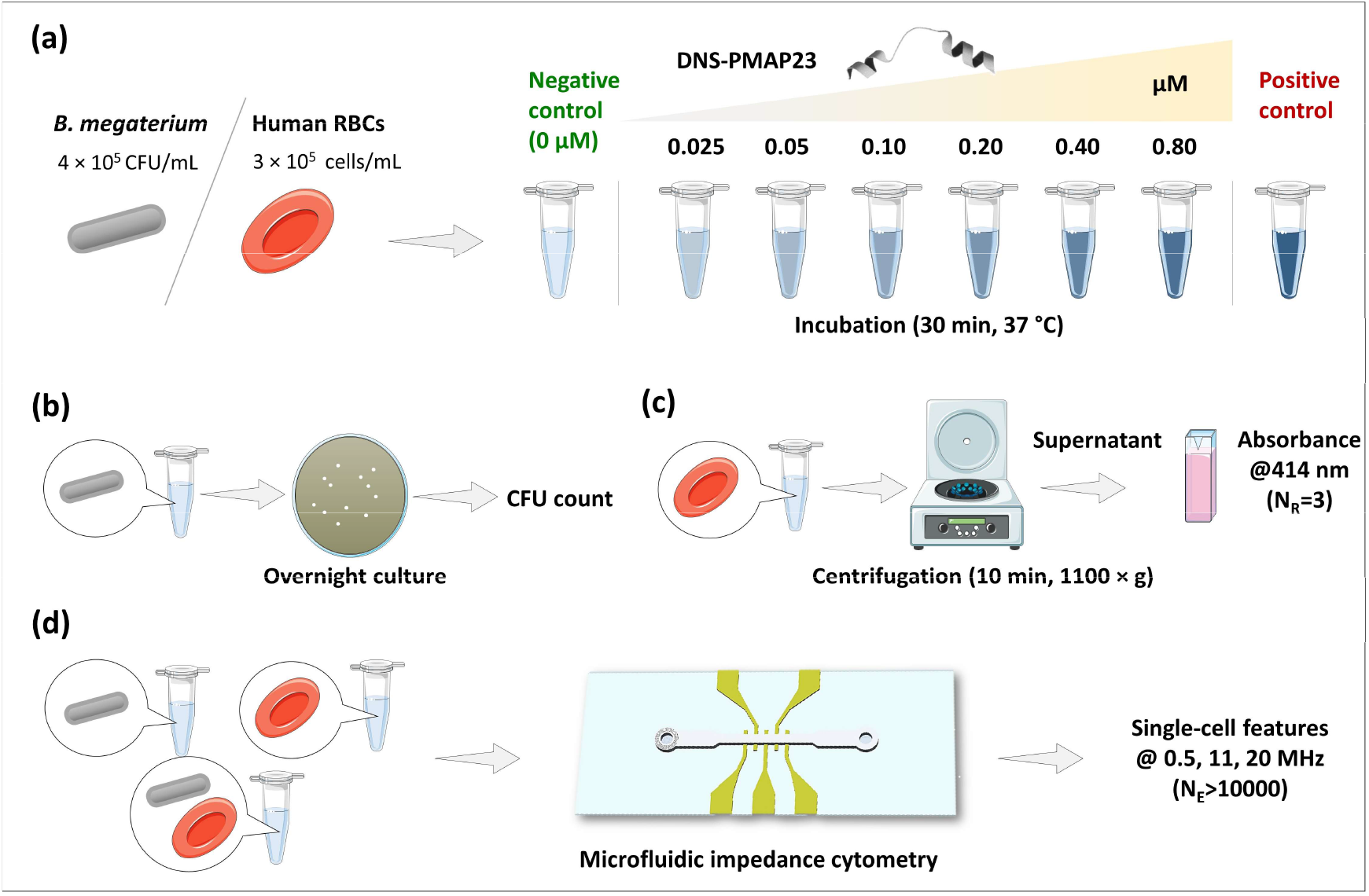
Overall experimental protocol. (a) Sample preparation: *B. megaterium* cells and human RBCs were incubated at 37 °C for 30 min with the DNS-PMAP23 peptide at six different concentrations in the range 0.025-0.80 μM. As negative control, samples without peptide were prepared. As positive control, samples with peptide concentration over 1 μM (for the bacteria) or samples treated with osmotic shock (for the RBCs) were prepared. (b) Standard bactericidal assay: colony-forming unit (CFU) count after overnight culture. (c) Standard RBCs hemolysis assay: absorbance measurements at 414 nm of supernatant after centrifugation (NR, number of replicates). (d) Microfluidic impedance cytometry: bacterial suspensions, RBCs suspensions, or mixed samples were measured with an impedance cytometer at 0.5 MHz, 11 MHz, and 20 MHz stimulation frequency. Thousands of single cells were acquired for each experimental condition (NE, number of events).

## 2. Materials and methods

### 2.1. Materials

DNS–PMAP23 (Dansyl-RIIDLLWRVRRPQKPKFVTVWVR-NH2), labeled with 5-(dimethylamino)naphthalene-1-sulfonyl (dansyl) at the N-terminus and amidated at the C-terminus, was purchased from AnyGen Co. (Gwangju, South Korea).

The Gram-positive bacterium *B. megaterium* Bm11 was kindly provided by Prof. Hans G. Boman (MTC, Karolinska Institute, Sweden). Red blood cells were collected from blood samples obtained from healthy volunteers.

### 2.2. Sample preparation

*B. megaterium* Bm11 was grown in LB (Luria−Bertani broth) medium at 37 °C in an orbital shaker until a mid-log phase was reached, as indicated by absorbance at 590 nm of 0.8. Bacterial cells were centrifuged (1400 × g for 10 min, Eppendorf 5702 centrifuge, Hamburg, Germany) and washed eight times in buffer A (5 mM HEPES, pH=7.3, 110 mM KCl, 15 mM glucose), to remove traces of LB medium. The cells were then resuspended in buffer A (Savini et al., 2020). Control experiments, performed in the absence of peptide, with approximately 3 × 10^7^ colony-forming unit (CFU) per mL, demonstrated that in this minimal medium, where bacteria are viable but do not multiply, the density of CFU remained constant, within experimental errors, for at least two hours, both at 25 °C and 37 °C, thus maintaining a constant density of live cells in the timeframe of the experiments. Bacterial cells were diluted in buffer A to a final cell density of 4 × 10^5^ CFU/mL and were incubated with DNS-PMAP23 at different concentrations (from 0.025 μM to 0.80 μM) at 37 °C for 30 minutes. Total bacterial killing (positive control) was obtained by using a high peptide concentration (> 1 μM). Negative control was obtained by suspending bacterial cells in buffer A, without any peptide. Bacterial samples were analyzed in parallel via a standard antibacterial activity assay and via microfluidic impedance cytometry.

Blood was washed six times with 5mM HEPES, pH 7.3, 150 mM NaCl (buffer E), and resuspended in the same buffer. After this step, RBC density was measured with an automated hematology analyzer Sysmex XE-2100 (TOA Medical Electronics, Kobe, Japan). Aliquots of the RBC suspension, diluted in buffer A at a final concentration of 3 × 10^5^ cells/mL, were incubated with six different concentrations of DNS-PMAP23 (from 0.025 μM to 0.80 μM), at 37 °C for 30 minutes. Negative control was obtained by suspending RBCs in buffer A, without any peptide. Total hemolysis (positive control) was obtained by suspending RBCs in distilled water overnight (osmotic shock). The RBC samples were analyzed in parallel via a standard hemolytic activity assay and via microfluidic impedance cytometry.

A mixed sample containing both bacterial cells and RBCs suspended in buffer A at concentrations of 4 × 10^5^ CFU/mL and 3 × 10^5^ cells/mL, respectively, was also prepared. An aliquot of this suspension was treated with peptide incubation (0.35 μM) at 37 °C for 30 minutes. The untreated and treated mixed samples were analyzed via microfluidic impedance cytometry.

### 2.3. Antibacterial activity assay

After peptide incubation, aliquots of 5 μL of bacterial cell suspension were withdrawn, diluted in buffer A to optimize the cell-density for colony counting, and spread onto LB-agar plates for counting after overnight incubation at 37 °C. Survival (*S*) of bacterial cells was expressed as fraction with respect to the untreated sample: *S* = *CFU*/*CFU*_*NC*_ (where the subscript *NC* denotes the negative control sample). The percentage of bacterial killing was calculated as follows:

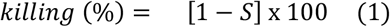

Datapoints were fitted with the Hill model:

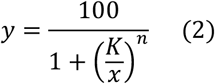

where *x* denotes the peptide concentration. The parameter *K* is the peptide concentration corresponding to half bacterial killing, whereas the parameter *n* is the Hill coefficient, which is an indicator of the cooperativity of peptide binding to cell.

### 2.4. Hemolytic activity assay

The hemolytic activity of DNS-PMAP23 was measured on human RBCs, following the previously published protocol (Savini et al., 2017). Briefly, after peptide incubation, the RBC samples were centrifuged in a Spectrafuge™ 24D (Labnet International Inc, Edison, NJ, U.S.A.) for 10 minutes at 1100 × g, and the absorbance (*Abs*) of the supernatant was measured on a Cary-UV 100 Scan spectrophotometer (Varian, Middelburg, Netherlands) at 414 nm (*i*.*e*., the wavelength of maximum absorbance of the Soret band) using 1 cm pathlength cuvettes.

The hemoglobin release (i.e., normalized absorbance) was calculated as follows:

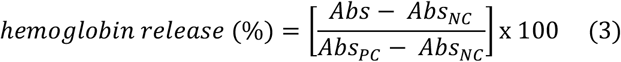

(where the subscript *PC* denotes the positive control sample). Datapoints were obtained as the mean of three independent measurements. The Hill model (**eq. 2**) was used to fit the datapoints.

### 2.5. Microfluidic impedance cytometry

The microfluidic impedance chip (**Fig. S1**, Supplementary Material) consisted of a PDMS fluidic layer sealed to a microscope glass slide (75 mm x 25 mm) patterned with microelectrodes (Ti/Au, 20 nm/200 nm). Standard techniques were used for device microfabrication, as previously described (Caselli et al., 2020). In the electrical sensing zone, channel width and channel height were 40 μm and 20 μm, respectively. Electrodes were 30 μm wide in the flow direction, with a 10 μm spacing. A three-electrode differential measurement scheme was used (**Fig. S1(a)**): AC voltage was applied to the central electrode; the currents flowing through the lateral electrodes were conditioned by a transimpedance amplifier (HF2TA, Zurich Instruments) and sent as input to an impedance spectroscope (HF2IS, Zurich Instruments); the spectroscope performed lock-in demodulation and the demodulated differential signals were saved into a PC for subsequent signal processing. Measurements at 0.5 MHz, 11 MHz, and 20 MHz stimulation frequency were performed.

Before the measurements, the samples were spiked with polystyrene beads (4.5 μm diameter, Polyscience) at a concentration of about 2 × 10^5^ beads per mL as an internal reference. A syringe pump (Harvard Apparatus) was used to inject the samples into the microfluidic chip (10 μL/min flow rate).

Thousands of single-cell events were measured for each experimental condition. Each measured event was fitted with a bipolar Gaussian template (cf. **Fig. S2**, Supplementary Material, for details). Electrical diameter (i.e., cube root of peak amplitude) and electrical phase were computed at each stimulation frequency. Beads signals were used to calibrate those features (i.e., the average electrical diameter of the bead population was set to 4.5 μm, and the average bead phase was set to zero), to enable quantitative comparison between measurements (Salahi et al., 2022).

The normalized phase of the bacterial population was calculated as follows:

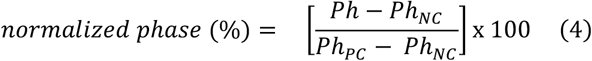

where *Ph* denotes the median phase. As detailed in **Section 3.2**, the impedance-based analysis of the RBCs revealed the presence of two subpopulations, labelled as subpopulation H and subpopulation L. Denoting by *f*L the relative fraction of RBCs belonging to subpopulation L, the normalized fraction of L-subpopulation was calculated as follows:

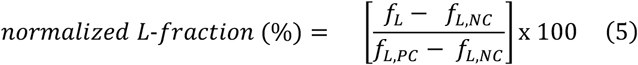

In analogy with the standard antibacterial and hemolytic activity assays, the Hill model (**eq. 2**) was used to fit the electrical signatures (namely, the normalized phase (eq. 4) and the normalized L-fraction (eq. 5)).

### 2.6. Image acquisition

With the purpose of controlling optically the passage of cells in the microchannel, the sample flow through the electrical sensing zone was acquired with a high-speed video microscopy system (Photron Mini UX100 camera operating at 4000 fps, 3.9 μs shutter time; Zeiss Axio Observer microscope with 20x objective), simultaneously to impedance acquisition, as described in previous works (D’Orazio et al., 2021; Reale et al., 2022).

## 3. Results

### 3.1. Impedance-based characterization of B. megaterium cells under peptide exposure

The results of the impedance-based characterization of *B. megaterium* cells exposed to the DNS-PMAP23 peptide are collected in **Figure 2. Figure 2(a)** shows the density plots of the phase against the electrical diameter, at each stimulation frequency (0.5, 11 and 20 MHz), for the negative control (i.e., 0 μM, no peptide), the sample at 0.10 μM and a sample at high peptide concentration (i.e., 2 μM). In the negative control sample (first column), at low frequency (0.5 MHz) the electric diameter falls in the range 2.5-3.5 μm and the phase is close to zero (i.e., phase of reference beads). Both the electrical diameter and the phase diminish by increasing the frequency from 0.5 MHz to 11 or 20 MHz. The effect of peptide incubation mainly affects the phase at high frequency. Specifically, at 11 MHz or 20 MHz the addition of the peptide induces a shift of the phase from negative values towards zero. On the other hand, at low frequency (0.5 MHz) the phase remains rather stable with peptide exposure. These trends are further visualized in **Figure 2(b)**, reporting the corresponding empirical probability density function of the phase at each frequency. The behaviour of the phase across the whole set of tested peptide concentrations (i.e., 0 μM, 0.025 μM, 0.05 μM, 0.10 μM, 0.20 μM, 0.40 μM, 0.80 μM, and 2 μM) is shown in **Figure 2(c)**, where the median phase values and the interquartile ranges are reported. The sensitivity to peptide exposure of the phase at high frequency (11 or 20 MHz) is confirmed.

**Fig. 2.**
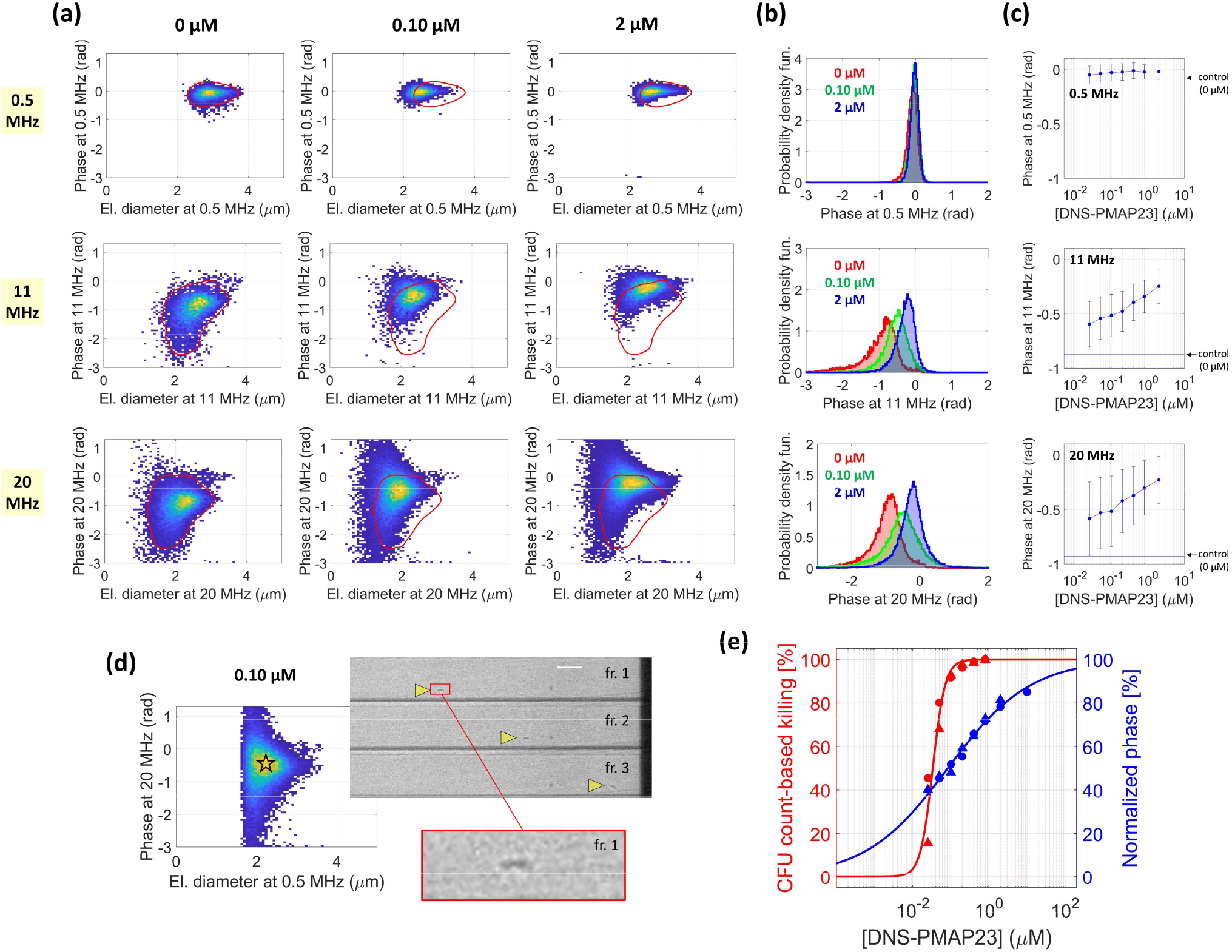
Results of the *B. megaterium* analysis. (a) Impedance-based characterization of *B. megaterium* cells at 0.5 MHz (first row), 11 MHz (second row), and 20 MHz (third row) stimulation frequency. The density plot of the phase against the electrical diameter is shown for the negative control sample (0 μM, first column), the sample at 0.10 μM (second column), and the sample at 2 μM (third column). For each stimulation frequency (i.e., in each row), the red contour line in the first column denotes the region enclosing 95% of the datapoints of the negative control sample (0 μM). This contour line is plotted as a reference also in the density plot of the samples at 0.10 μM (second column) and 2 μM (third column). (b) Empirical probability density function of the phase at 0.5 MHz (first row), 11 MHz (second row), and 20 MHz (third row), for the samples at 0 μM (in red), 0.10 μM (in green), and 2 μM (in blue). (c) Median values of the phase at 0.5 MHz (first row), 11 MHz (second row), and 20 MHz (third row) as a function of the peptide concentration. Interquartile ranges are also shown. In each panel, the horizontal line indicates the median value of the phase of the negative control sample (0 μM). (d) Density plot of the phase at 20 MHz against the electrical diameter at 0.5 MHz (sample at 0.10 μM), along with exemplary snapshots of a flowing *B. megaterium* cell (three consecutive frames, fr.; scale bar is 20 μm). (e) Comparison of the killing curve based on microfluidic impedance cytometry (at 20 MHz), in blue, with the killing curve based on CFU counts, in red. Markers denote experimental datapoints (circles and triangles refer to different experiment repetitions), continuous lines denote fits of the results.

By using the impedance signals as pointers to image frames (D’Orazio et al., 2021), bacterial cells could be automatically identified in the acquired high-speed videos. Whereas the electrical signatures of the bacteria were sensitive to peptide exposure, no noticeable differences between unexposed and exposed bacteria were found at optical inspection. Exemplary snapshots of an individual bacterial cell flowing through the microfluidic cytometer are shown in **Figure 2(d)**.

The standard assay for determining peptide bactericidal activity is based on CFU count after overnight culture. The corresponding bacterial killing (**eq. 1**) at different peptide concentrations is reported in red in **Figure 2(e)**. The relevant fit (**eq. 2**) is also shown (parameter values, mean ± std: *K* = 0.034 ± 0.003 μM, *n* = 2.7 ± 0.6). The CFU-count based method shows that *B. megaterium* is susceptible to the DNS-PMAP23 peptide, since the number of bacterial cells able to form colonies diminishes for increasing peptide concentration. The phase at high frequency is sensitive to peptide exposure, hence it is a potential biomarker of peptide bactericidal activity that does not require bacterial cultures. The normalized phase (**eq. 4**) at the different peptide concentrations is reported in blue in **Figure 2(e)**, along with the corresponding fit (parameter values: *K* = 0.08 ± 0.03 μM, *n* = 0.40 ± 0.06). The bacterial killing caused by the treatment with DNS-PMAP23 at increasing concentration is reflected in an increase of the normalized phase, even though the fitted parameters turned out to be different.

### 3.2. Impedance-based characterization of RBCs under peptide exposure

The results of the impedance-based characterization of RBCs exposed to the DNS-PMAP23 peptide are collected in **Figure 3**. As a representative example, **Figure 3(a)** shows the density plots of the phase against the electrical diameter, at each stimulation frequency (0.5, 11 and 20 MHz), for the negative control (i.e., 0 μM, no peptide), the samples incubated at different peptide concentrations (0.025 μM, 0.05 μM, 0.10 μM, 0.20 μM, 0.40 μM, 0.80 μM) and the positive control (osmotic shock). At low frequency (0.5 MHz) the electrical features do not exhibit a noticeable trend with respect to peptide exposure. For all samples, the electrical diameter at 0.5 MHz falls in the range 4.5-7.5 μm and the phase is slightly lower than that of the reference beads (i.e., around -0.2 rad). At high frequency (11 or 20 MHz), lower electrical diameters are measured (2.5-6 μm range) and a distinctive behaviour is found, characterized by the presence of two RBCs subpopulations having different phases. The subpopulation with higher phase is denoted by H, the subpopulation with lower phase is indicated by L. Subpopulation L also exhibits a noticeable reduction of the electrical diameter at 20 MHz compared to that at 11 MHz. The relative fraction of the two subpopulations varies across the samples. In the negative control sample (0 μM) and at low peptide concentration (up to 0.10 μM in this representative example), subpopulation H is markedly dominant. At 0.20 μM both subpopulations are well represented, whereas at 0.40 μM and 0.80 μM subpopulation L is markedly dominant. The osmotic sample exhibits one population, which has electrical signatures close to those of subpopulation L (even though they are not completely overlapping). Interestingly, subpopulations H and L, despite having comparable electrical size (i.e., electrical diameter at 0.5 MHz), turned out to be remarkably different at optical inspection (**Figure 3(b)**). Whereas cells belonging to subpopulation H were clearly identifiable in the recorded high-speed videos, cells belonging to subpopulation L were invisible with the present optical setup. Cells of the osmotic sample were not optically detectable, too.

**Fig. 3.**
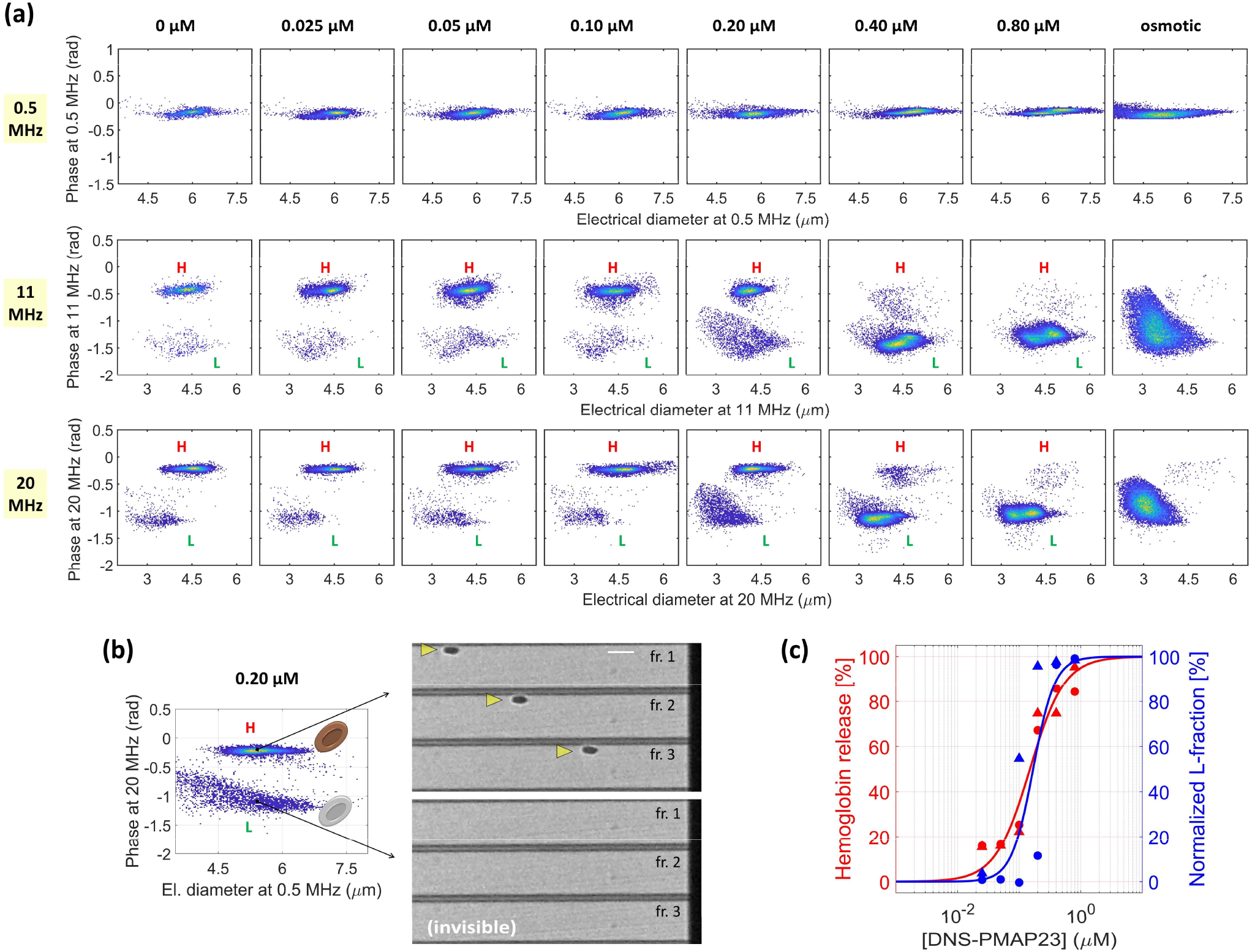
Results of the RBC analysis. (a) Impedance-based characterization of RBCs at 0.5 MHz (first row), 11 MHz (second row), and 20 MHz (third row). For each peptide concentration (column-by-column), the density plot of the phase against the electrical diameter is shown (last column refers to the osmotic sample). Two RBCs populations are found at 11 MHz and 20 MHz, denoted by H (high phase) and L (low phase). (b) Density plot of the phase at 20 MHz against the electrical diameter at 0.5 MHz (sample at 0.20 μM). RBCs of subpopulation H are clearly detectable in the high-speed video (three consecutive frames, fr.; scale bar is 20 μm). RBCs of subpopulation L, despite having comparable electrical size, resulted invisible at optical inspection. (c) Comparison of the RBC toxicity assay based on microfluidic impedance cytometry (at 20 MHz), in blue, with the standard toxicity assay based on hemoglobin release, in red. Markers denote experimental datapoints (circles and triangles refer to RBCs from two different donors), continuous lines denote fits of the results.

The standard assay for determining peptide toxicity to RBCs is based on the quantification of hemoglobin release, as measured by absorbance levels in a sample where RBCs have been removed by centrifugation. **Figure 3(c)** shows, in red, the hemoglobin release (**eq. 3**) for the tested peptide concentrations along with the relevant fit (parameter values: *K* = 0.15 ± 0.02 μM, *n* = 1.5 ± 0.3). The results indicate that RBCs are susceptible to the DNS-PMAP23 peptide, in agreement with previous reports (Savini et al., 2017). **Figure 3(c)** also shows the normalized fraction of subpopulation L (**eq. 5**) and the corresponding fit (parameter values: *K* = 0.17 ± 0.04 μM, *n* = 2.5 ± 1.4). The hemoglobin release caused by the incubation of RBCs with DNS-PMAP23 at increasing concentrations is reflected in a progressive increase of subpopulation L, with similar values of the fitted parameters. These results suggest that the latter subpopulation represents damaged RBCs, whereas subpopulation H, which is dominant at low peptide concentration, represents healthy RBCs.

### 3.3. Characterization of a mixed sample of B. megaterium cells and RBCs

**Figure 4(a)** shows the density plot of the electrical diameter at 0.5 MHz against the electrical diameter at 20 MHz for the untreated (i.e., no peptide incubation) mixed sample. Three subpopulations are found. Based on the analysis of the separate samples (**Figures 2(a)** and **3(a)**), the subpopulation with lower electrical diameter at 0.5 MHz (i.e., < 3.8 μm) corresponds to bacterial cells, while the two subpopulations with higher electrical diameter at 0.5 MHz (i.e., > 3.8 μm) correspond to RBCs. This criterion has been used to study the two cell types separately in subsequent analyses. **Figure 4(b)** and **(c)** show the density plot of the phase at 20 MHz against the electrical diameter at 20 MHz for the bacterial cells and for the RBCs, respectively. Of the two RBC subpopulations, the main one shows higher values of both the electrical diameter and the phase (i.e., -0.2 rad median phase value against -0.9 rad). In agreement with the analysis of the RBC sample reported in **Section 3.2**, the main RBC subpopulation corresponds to healthy RBCs (subpopulation H), while the minor subpopulation represents damaged RBCs (subpopulation L). **Figure 4(d)** shows the empirical probability density function of the phase at 20 MHz for the three subpopulations.

**Fig. 4.**
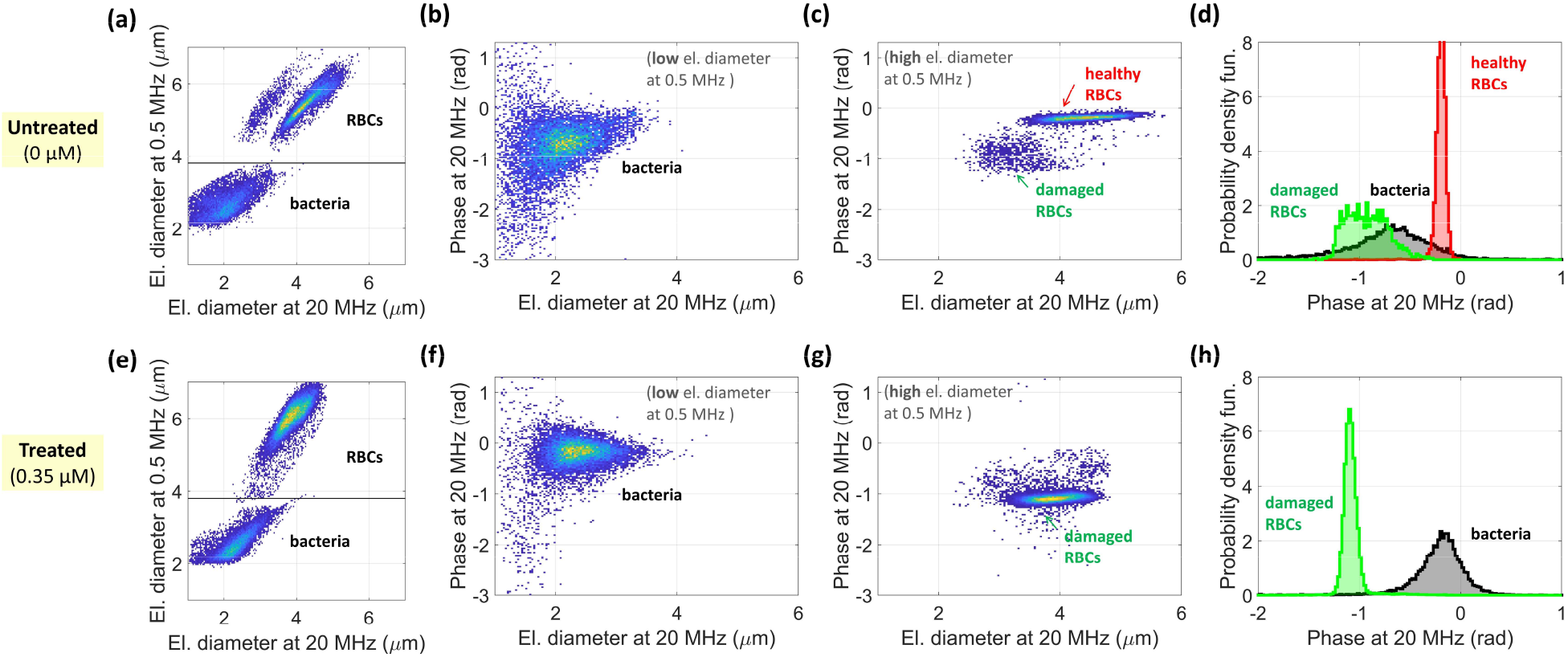
Results of impedance-based analysis of mixed samples (i.e., containing both *B. megaterium* cells and RBCs): (a)-(d) untreated sample (0 μM), (e)-(h) treated sample (0.35 μM). (a) and (e) Density plot of the electrical diameter at 0.5 MHz against the electrical diameter at 20 MHz, with highlight of relevant subpopulations. The gating line is also shown (electrical diameter at 0.5 MHz equal to 3.8 μm). (b) and (f) [resp. (c) and (g)] Density plot of the phase at 20 MHz against the electrical diameter at 20 MHz for the bacterial cells [resp. for the RBCs]. (d) and (h) Empirical probability density function of the phase at 20 MHz, for each subpopulation.

The analysis of the treated (i.e., peptide incubation at 0.35 μM) mixed sample is reported in **Figure 4(e)-(h)**. Two subpopulations appear in the density plot of the electrical diameter at 0.5 MHz against the electrical diameter at 20 MHz (**Figure 4(e)**), which are identified as bacteria (electrical diameter at 0.5 MHz < 3.8 μm) and RBCs (electrical diameter at 0.5 MHz > 3.8 μm). The separate density plots of the phase at 20 MHz against the electrical diameter at 20 MHz for the bacterial cells and for the RBCs are shown in **Figure 4(f)** and **Figure 4(g)**, respectively, while the empirical probability density function of the phase at 20 MHz is reported in **Figure 4(h)**, for both subpopulations. Peptide exposure induces an increase in the median phase of bacterial cells (i.e., from -0.7 to -0.2 rad). A similar trend was obtained with the sample containing bacterial cells alone. Furthermore, the RBC subpopulation in the treated sample exhibits the reduced phase typical of the damaged RBCs.

## 4. Discussion

AMPs hold promises to fight AMR. However, AMP development requires AST methods that are fast (minutes) and able to assess AMP activity and cytotoxicity in presence of both bacterial and eukaryotic cell types, with single-cell sensitivity. Among the microfluidic technologies that are being explored in AST research (Postek et al., 2022; Qin et al., 2021), single-cell impedance cytometry seems particularly suited to meet these requirements (Honrado et al., 2021b).

Impedance cytometry provides single-cell electrical fingerprints that convey information about intrinsic cell properties. At low frequency, cells with intact membrane/cell-wall behave as insulating particles and the impedance signal is proportional to cell volume. Changes in membrane/cell-wall capacitance are observed in the mid-frequency range, whereas changes in cytoplasmic properties affect the high frequency part of the spectrum (Gawad et al., 2004; Honrado et al., 2021b; Spencer et al., 2020). Therefore, the effects of AMPs on both bacterial and host cells were predicted to produce measurable changes in impedance values, since they include perturbation of cell membrane permeability, dissipation of transmembrane gradients, loss of intracellular material, and changes in cell volume/shape (Hartmann et al., 2010; Marcellini et al., 2010; Matsuzaki, 2019; Semeraro et al., 2022). In addition, in the case of bacteria, after membrane perturbation, AMPs accumulate inside dead cells (Kaji et al., 2021; Semeraro et al., 2022; Snoussi et al., 2018; Wu and Tan, 2019), binding to intracellular components (Savini et al., 2020) and causing rigidification of the cytosol (Zhu et al., 2021, 2019).

In this work, the frequency-dependent cell electrical phenotypes were quantified in terms of the electrical diameter and the electrical phase, which are common metrics for impedance-based cell characterization. The electrical diameter was recently used to characterize the response of susceptible bacteria to traditional antibiotics. For β-lactam type antibiotics, which mainly work by inhibiting cell wall biosynthesis, either an increase or a reduction of the low-frequency electrical diameter was reported, depending on the bacterial type (Gram negative *vs* Gram positive) (Spencer et al., 2020; Tang et al., 2022). For colistin, which affects membrane permeability in Gram-negative bacteria, a reduction in the electrical diameter of *Klebsiella pneumoniae* was found (Spencer et al., 2020). Broadly speaking, the mode of action of DNS-PMAP23 (membrane perturbation) is analogous to that of colistin, even though the latter targets lipopolysaccharides (Sabnis et al., 2021), while the former does not interact with them specifically (Savini et al., 2020). Our results (**Figure 2(a)**) did not show specific alterations of the electrical diameter of *B. megaterium* (Gram positive) upon exposure to different peptide concentrations. This may be due to the different structure of Gram-positive compared to Gram-negative bacteria and/or to the short time-window of the measurement (within which any change in volume may not be fully developed). On the other hand, upon peptide exposure we found significant variations of the high-frequency phase of *B. megaterium* (**Figure 2(b)** and **(c)**). Specifically, the phases at 11 MHz and 20 MHz increase with increasing peptide concentration. A similar increase in the high-frequency (40 MHz) phase was recently reported for *Staphilococcus aureus* (Gram positive) exposed to Cefoxitin (β-lactam type antibiotic) (Spencer et al., 2020). The electrical phase was also used in the literature to monitor bacteria inactivation (Bertelsen et al., 2020; David et al., 2012) and bacterial spore germination (Moore et al., 2020).

The impedance-based characterization of the RBCs revealed the presence of two subpopulations (**Figure 3(a)** and **(b)**), which were identified as healthy RBCs and damaged RBCs. Their electrical diameters were similar irrespective of frequency (in fact, the electrical diameter of the damaged RBC subpopulation was slightly smaller than that of the healthy subpopulation at 20 MHz). Their phases were similar at 0.5 MHz (and slightly lower than that of the reference beads), whereas the phase of the damaged subpopulation showed a significant reduction at 11 MHz and 20 MHz. By referring to the single-shell model - which is commonly used for the interpretation of the RBC impedance spectrum (Salahi et al., 2022; Spencer et al., 2020) - the behaviour of damaged RBCs compared to healthy RBCs is compatible with an increase in the membrane conductivity (due to pore formation) and an increase of the intracellular conductivity and permittivity (due to hemoglobin release and cytoplasm replacement by suspension medium) (cf. **Figure S3** of the Supplementary Material). These structural modifications are reflected in an altered optical behaviour (damaged RBCs turned out to be invisible with the present imaging setup).

Based on the electrical characterization, we were able to build the DNS-PMAP23 antibacterial activity and RBC cytotoxicity curves. A general agreement with the activity and cytotoxicity curves obtained with the reference methods was found (**Figures 2(e)** and **3(c)**). The RBC cytotoxicity assays based on impedance or hemoglobin release showed comparable behaviours (**Figure 3(c)**). Regarding the antibacterial activity, the Hill coefficient provided by the standard approach was noticeably higher than that of the impedance-based approach (i.e., 2.7 vs 0.4). A possible explanation for this observation is that the two approaches are measuring different effects. In fact, the standard assay accounts for biological cell changes (bacterial killing), whereas the impedance-based approach accounts for biophysical cell changes (structural modifications). Since, as discussed above, high frequency phase is influenced mainly by the cytosolic properties, the impedance signal could reflect peptide accumulation inside the killed bacterial cells, which is known to take place following membrane perturbation (Kaji et al., 2021; Savini et al., 2020; Semeraro et al., 2022; Snoussi et al., 2018; Wu and Tan, 2019), and might increase further if AMP concentrations higher than those required for membrane perturbation are added. It is currently debated whether peptide accumulation inside the cytosol is simply a consequence of membrane disruption, or if it is required for bacterial killing (Savini et al., 2020; Semeraro et al., 2022). The difference observed between the antibacterial curves obtained by measuring CFUs and by impedance cytometry might support the former hypothesis. This aspect is currently being investigated further.

The impedance measurements were performed right after peptide incubation, without any additional preparation step. For each experimental condition, several thousands of cells were measured at a throughput of about a hundred cells per second (∼5 min acquisition time). Compared to the long times (overnight incubation) required by the CFU-count based activity assay, the rapidity of the impedance-based assay is a major advantage. The timeframe of the standard cytotoxicity assay based on hemoglobin release is not critical, since it is essentially set by the 10 min centrifugation step, with the possibility of parallel sample processing. However, the standard cytotoxicity assay lacks single-cell sensitivity and cannot provide information on sample heterogeneity (i.e., the presence of RBC subpopulations).

Whereas substantially different assays are traditionally used for activity testing (i.e., CFU-count based bacterial killing) and for cytotoxicity testing (i.e., absorbance-based RBCs hemoglobin release), microfluidic impedance cytometry can serve both purposes. The technique also has the potential for simultaneous AST of bacteria and determination of toxicity to host cells, as shown by the reported proof-of-concept experiment with a mixture of *B. megaterium* and RBCs (**Figure 4**). A sample where host cells and pathogens coexist mimics an infection in vivo significantly better than the separate assays normally employed to assess AMP selectivity (Loffredo et al., 2021; Savini et al., 2017). In view of its rapidity and versatility, impedance cytometry is uniquely posed to develop next-generation AST approaches and holds promises for AMPs screening and optimization.

## 5. Conclusion

In this work, we presented the application of single-cell microfluidic impedance cytometry to assess the susceptibility of bacterial cells (*B. megaterium* cells) and host cells (RBCs) to a representative antimicrobial peptide (DNS-PMAP23). Impedance cytometry turned out to be an effective way for rapid assessment of both antimicrobial activity and RBC cytotoxicity, as confirmed by comparison with standard bacterial killing assays and hemolytic activity assays. Overall, the main merits of the proposed technique with respect to traditional approaches are: (i) the suitability to both bacterial and host cells (even in the same sample), (ii) the rapidity of the analysis (thousands of cells measured in a few minutes), (iii) the single-cell sensitivity (which enables subpopulations analysis). Further studies will focus on the characterization of mixed samples with bacteria and RBCs at different relative concentrations, to shed light into possible interactions and peptide sequestration mechanisms.

## Acknowledgments

This work was supported by the Regione Lazio (Research Groups 2020 Programme, E85F21002390002, to F.C. and A.D.N.) and by the Italian Ministry of University and Research (PRIN 2020833Y75, to L.S.), and by the National Research Foundation (NRF) of Korea (No. NRF-2022M3A9H5096106, to Y.K.P.).

## Author contributions

**Conceptualization**: L.S., F.C., P.B. **Methodology**: L.S., F.C., P.B., M.L.M., C.T., A.D.N., B.C. **Software**: F.C., P.B. **Formal analysis**: F.C., C.T., L.S., P.B. **Investigation**: C.T., A.D.N., F.C., B.C., R.M., F.R. **Resources**: A.D.N., F.C., L.S., Y.K.P., M.L.M., R.M. **Writing - Original Draft**: F.C., L.S., C.T. **Writing - Review & Editing**: all Authors **Visualization**: F.C., C.T., B.C., L.S. **Supervision**: L.S., M.L.M. **Project administration**: F.C., L.S. **Funding acquisition**: L.S., F.C., A.D.N., Y.K.P.

