## Supplementary material for "Rapid assessment of susceptibility of bacteria and erythrocytes to antimicrobial peptides by single-cell impedance cytometry"

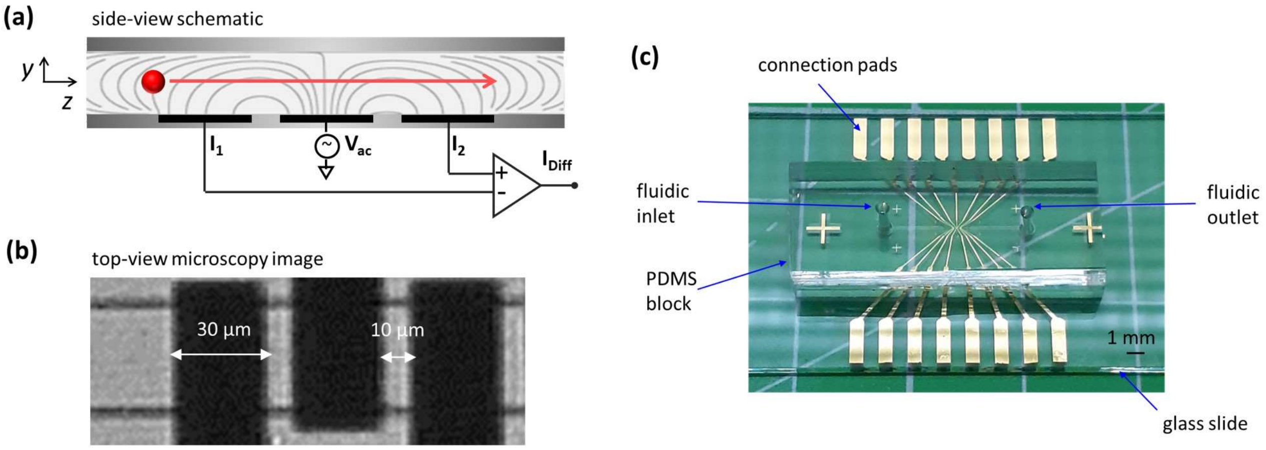


**Fig. S1.** (a) Schematic representation of the coplanar-electrode microfluidic impedance cytometer (side view): AC excitation signals ($V_{ac}$) at different stimulation frequencies are simultaneously applied to the central electrode, and the difference in current flowing through the lateral electrodes is measured, $I_{Diff} = I_2 - I_1$. (b) Microscopy image of the electrical sensing zone (top view). Dark bands denote electrodes. Relevant dimensions are indicated. (c) Image of the microfluidic impedance chip. The chip is made of a PDMS block containing the microfluidic channel, bonded to a glass slide with deposited Ti/Au microelectrodes. The pads for the electric connections and the fluidic access ports are indicated in the picture.

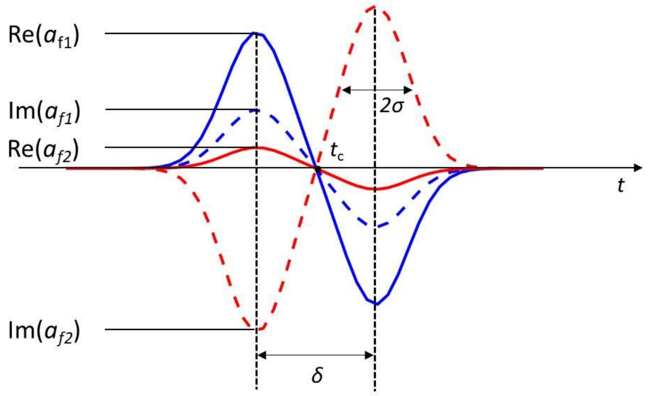


**Fig. S2**. Bipolar Gaussian template used to fit the single-cell events:

$$s(t)_f = a_f\left(e^{-\frac{(t-(t_c-\delta/2))^2}{2\sigma^2}} - e^{-\frac{(t-(t_c+\delta/2))^2}{2\sigma^2}}\right) \qquad (1)$$

The template is characterized by the complex frequency-dependent amplitude $a_f$, the peak-width control $\sigma$, the peak-to-peak time $\delta$ and the central time $t_c$. The complex amplitude $a_f$ is converted in modulus and phase. Since at low frequencies signal amplitude is proportional to cell volume, its cube root provides an electrical measure of the cell diameter (the electrical diameter).

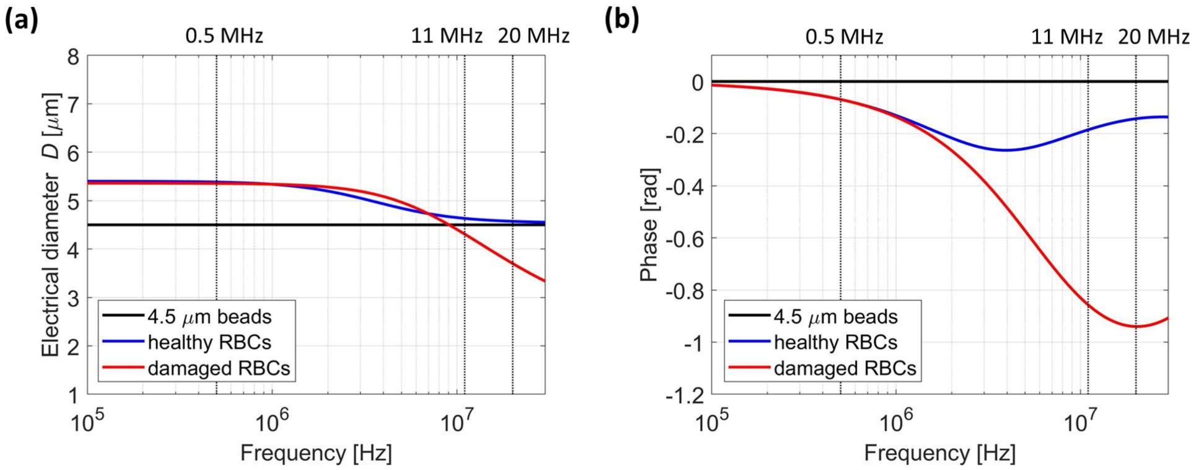


**Fig. S3**. Simulated impedance spectra for beads, healthy RBCs, and damaged RBCs, based on Maxwell's mixture theory and the single-shell model (cf. e.g. [1], [2] for the relevant theory). The three experimental frequencies (0.5 MHz, 11 MHz, and 20 MHz) are highlighted. Simulation parameters are collected in **Table S1** below. Beads and healthy RBCs parameter values are taken form Refs. [2], [3]. Buffer conductivity and relative permittivity are relevant to PBS. To mimick peptide-induced pore formation, the damaged RBCs have higher membrane conductance than healthy RBCs. Moreover, damaged RBCs interior parameters are set assuming that 80% of the cytoplasm is replaced by the buffer. This model explains the lower high-frequency phase (and electrical diameter at 20 MHz) exhibited by damaged RBCs with respect to healthy RBCs.

**Table S1**. Simulation parameters.

| **Particle/medium** | **Diameter [μm]** | **Interior conductivity [S/m]** | **Interior relative permittivity [-]** | **Membrane capacitance [mF/m²]** | **Membrane conductance [S/m²]** |
|---|---|---|---|---|---|
| 4.5 μm beads | 4.5 | $2.7 \times 10^{-3}$ | 2.5 | - | - |
| Healthy RBCs | 5.4 | 0.52 | 65 | 8.89 | $10^3$ |
| Damaged RBCs | 5.4 | 1.4 | 77 | 8.89 | $10^4$ |
| Buffer | - | 1.6 | 80 | - | - |
